# Mating system modulates mate selection in aphidophagous ladybird, *Menochilus sexmaculatus*

**DOI:** 10.1101/2020.03.19.991737

**Authors:** Swati Saxena, Geetanjali Mishra, Omkar

## Abstract

Mate competition and selection of mates is strongly influenced by the organism’s mating system. Monogamous matings provide more benefits as compared to polygamous matings. However, it has been proven that in polygamous systems, females gain benefits from the males, therefore indulging in multiple mating, leading males to access maximum females. In zigzag ladybird, *Menochilus sexmaculatus*, mate choice has been studied on several pre-and postcopulatory factors. However, mate choice as a function of mating system is still remains untouched. In the present study, we hypothesised that the mating system modulates mate selection of beetles. Adults were held in different mating systems and then males and females were tested in mate choice trials. Polygynous males were more preferred over monogynous males. However, males preferred monandrous females over polyandrous females. In a second experiment, we also included relatedness as additional factor. In female mate choice trials, females preferred unrelated monogynous males which were earlier rejected over related polygynous and in male mate choice trials, males preferred unrelated polyandrous females over related monandrous females. The results were not confined only to mate choice but significant effect was also observed on time to commence mating, copulation duration, fecundity and percent egg viability.

## INTRODUCTION

Mating systems describe how pairing of male and female occurs while choosing a mate. It basically consists of two types of mating systems *i*.*e*. monogamy and polygamy. Monogamy involves the mating with a single mate (repetitive matings). In case of females, it can be monandry and in case of males it is called mongyny. If both the sexes mate with multiple mates they are considered polygamous (Emlen & Oring, 1977; Thornhill & Alcock, 1983; Davies, 1991; Shuster & Wade, 2003). While monogamy is much prized in the human social context, true genetic monogamy is extremely rare in the animal world, and even in presumably monogamous systems, single matings in a lifetime are rarest of the rare cases. In spite of several costs of multiple matings, such as reduced foraging, reduced longevity or even risks of physical injury such as damage of the female’s reproductive tract (Thornhill & Alcock, 1983; Arnqvist, 1989; Yasui, 1998; Arnqvist & Nilsson, 2000; Rolff & Siva-Jothy, 2002; Simmons, 2005), multiple matings are common to gain reproductive fitness. For female mating with different males is termed as polyandry and for males mating with different females called as polygyny. Some researchers have proposed that mating system reflects how much parental care each sex provides (Vehrencamp & Bradbury, 1984; Reynolds, 1996; Shuster & Wade, 2003). There are several theories, such as Bateman’s principles (Bateman, 1948), parental investment (Trivers, 1972), etc. that demonstrate that males produce numerous, small gametes (sperm) while females produce large and limited number of gametes (eggs). Experiments on *Drosophila* by Bateman revealed that males gain reproductive success by multiple matings. However, these predictions were not in accordance with the empirical data. Several researchers have suggested that females also mate with different males to gain benefits (Tang-Martinez, 2010; Gowaty et al.,2012). A number of hypotheses have been proposed to explain why polyandry or multiple matings exists in spite of it being costly. In many species, such as in *Drosophila subobscura* Collin, females are monogamous but upon receipt of certain nuptial gifts they mate multiply (Smith, 1956; Holman et al., 2008). In jewel wasps, *Nasonia vitripennis*, wild females are generally monogamous and laboratory females are reluctant to remate (Burton-Chellew et al., 2007). However, after females were reared for longer periods under laboratory conditions, they tend to mate multiply. Studies in *N. vitripennis* also revealed that tendency to mate multiple times has been found to be heritable (Shuker et al., 2007).

Multiple matings have their costs and benefits (Thornhill &Alcock, 1983). In the seaweed fly, *Coelopa frigida* Fabricius, (Blyth & Gilburn, 2006), females resist matings with mounted males by shaking, kicking, curving which are energetically costly responses. In water striders, mating is costly to females by increasing their probability of predation or decreasing their longevity due to physical damage (Arnqvist, 1989). Besides physical injury, chemical damage has also been reported in *D. melanogaster*. Male ejaculate may contain some toxic substances that can cause side effects on the female life span (Gems & Riddle, 1996; Siva-Jothy et al.,1998; Fowler & Partridge, 1989; Chapman et al., 1995), and decrease remating rate and sexual receptivity (Eberhard, 1996). In certain instances, it was found that multiple matings hamper the reproductive success (Ronn et al., 2006; Bezzerides et al., 2008; Pai & Beenasconi, 2008). In the cabbage beetle, *Colaphellus bowringi* Baly, monogamy was more advantageous for female than polyandry (Liu et al., 2010). However, contrary to this monogamy increase the chances of homozygosity. This was found to affect the social interactions of offspring, with those born to monogamous pairs having less prominent interactions than those born to the heterozygous offspring of polygamous pairs (Evans & Kelly, 2008). Besides all these costs, there are several benefits of multiple matings. They can be categorised into non-genetic or direct benefits and genetic or indirect benefits. Several empirical studies have addressed the effects of multiple mating on female fitness in various ways (Ridley, 1988; Vahed, 1998) focusing on the act of mating per se, the presence of sperm or the transfer of accessory substances. A large number of accessory substances are transferred to females that enhance the female reproductive performance (Gromko et al., 1984) and also affect the egg production (Opp & Prokopy, 1986) and laying. Indirect benefits include offspring attractiveness (‘sexy-sons’ hypotheses), their viability (Tregenza & Wedell, 2002; Gowaty et al.,2010; Omkar et al.,2010; Firman and Simmons, 2012), genetic heterogeneity, and phenotypic diversity (Barbosa & Magurran, 2010; Foerster et al., 2003).

Ladybirds are considered highly promiscuous; they can copulate for long time and change their mating partners frequently (Nedved & Honek, 2012). Polymorphism in ladybirds also tends to be one of the reasons behind multiple matings (Slogget & Honek et al.,2012) as it is one of the genetic marker and it is often maintained as a result of sexual selection (Muggleton et al.,1975; O’Donald & Majerus,1984). Omkar et al. (2010) reported that 11 matings were required to maximize the fecundity in *Coelophora saucia* (Mulsant). Studies on *Coccinella transversalis* F. and *C. septempunctata* reported that male virility diminishes with increase in number of matings, thus reducing female’s fecundity (Michaud et al., 2013). In *Menochilus sexmaculatus* (Fabricius) a positive effect of multiple mating on fecundity was recorded (Dubey, 2016).

Thus, in contention of the above information, it was hypothesised that polygamous individuals will be more preferred over monogamous in *M. sexmaculatus*. The present study was conducted to observe the mate choice between individuals of different mating system by monogamous and polygamous males or females. As it was earlier studied that these beetles show rejection towards related individuals (Saxena et al., 2016), to extend our study the choice was also given between related and unrelated individuals and individuals of different mating systems to study whether mating system can alter the rejection behaviour of these beetles towards related partners.

## MATERIALS AND METHODS

### Experimental species

*Menochilus sexmaculatus*, commonly known as zigzag beetle, was selected for experimentation due to its abundance in local fields, high reproductive output, and wide prey range (Agarwala & Yasuda 2000). Presence of basic information about pre-(Dubey et al., 2016a, b) and post-copulatory mate choice behaviour (Chaudhary et al., 2015) further provides a good foundation to build upon.

### Collection and rearing of experimental species

Adults of *M. sexmaculatus* (50 beetles) were collected from the agricultural fields of Lucknow, India (26°50’N, 80°54’E). Adult males and females were paired in plastic Petri dishes (hereafter, 9.0 × 2.0 cm) and were fed on *ad libitum* supply of cowpea aphid, *Aphis craccivora* Koch (Hemiptera: Aphididae) raised on cowpea, *Vigna unguiculata* L. reared in a glasshouse (at 25±2°C, 65±5% R.H.). The Petri dishes containing mating pairs were placed in Biochemical Oxygen Demand (BOD) incubators (Yorco Super Deluxe, YSI-440, New Delhi, India) at 27±1°C; 65±5% R.H.; 14L:10 D and inspected twice daily (1000 and 1500h) for oviposition. The eggs laid were separated daily and held in plastic Petri dishes (size as above) until hatching. Each hatched larva was reared individually in Petri dishes (size as above) until adult emergence.

### Rearing of adults of different mating system

Monogamous adults were prepared from the stock culture. For monandrous females, 5-day old unmated females were allowed to mate with 5-day-old unmated males. After their natural dislodging, both male and female were kept separately in new Petri dishes (size as above) and provided *ad libitum* supply of aphids over a period of 5 days. Thereafter, the same pair was reformed and allowed to mate at 24 hours interval, leading to the total of five matings.

For polyandrous females, 5-day old unmated female was allowed to mate with 5-day old unmated unrelated male until the completion of the mating. After postmating, beetles were separated. After 24 hours same females were provided with new once-mated male. Mating with different males continued for the next 5-days (mating status of the new males were kept same as that of female). Thus, monogynous 5-day-old males were allowed to mate with same unmated 5-day-old unrelated females at the interval of 24 hours for the next 5 days, whereas polygynous condition, 5-day old unmated males were allowed to mate with different females of the same mating status at every 24 hours for the next five days.

### Effect of mating system and relatedness on mate choice

#### (a) Mating system effects

In female mate choice trials, 10-day old monogynous and polygynous males were introduced simultaneously in the new Petri dish (size as above). A focal 10-day-old monandrous female was placed into this arena within 5 minutes of introducing the males. This setup allowed full interaction of the focal monandrous female with both the males present in the arena. If mating occurred, the unselected male was removed and mating was allowed to complete. If no choice was made within 30 minutes, the trial was discarded. To distinguish between different males, the adults were colour marked with a yellow or green dot on the posterior edge of elytra. To avoid the biasness due to colour marks, it was randomized in all the replicates of the mate choice trials. Similar mate choice trials were conducted for polyandrous females. All adults were unrelated in these treatments.

In male mate choice trials, one 10-day-old monogynous male was introduced in the new Petri dish (size as above). 10-day old unrelated monandrous and polyandrous females were introduced simultaneously. If no choice was made within 30 minutes, the replicate was discarded. Marking of both the females was done in similar way as above. Similar mate choice trials were conducted for polygamous males. All the mate choice trials were conducted in 15 replicates.

#### (b) Relatedness and mating system effects

To additionally study the effect of relatedness and the mating system, newly emerged females (0-2h old) were taken from the stock of related individuals, which was created and maintained as per Saxena et al. (2016), where the offspring of the random pairings from a laboratory stock populations were mated with their siblings creating a related stock. A 5-day old related female (female from the related stock) was allowed to mate with one related males for the next five days at 24 hours interval. Such females were called as related monogamous females. For related polyandrous females, they were allowed to mate with different related males for next five days at 24 hours interval. Status of the male was kept according to that of females.

Similarly, a 5-day-old male taken from the above related stock was allowed to mate with single related female for the next five days at 24hours intervals (related monogynous males). Another 5-day old related male was allowed to mate to with different related females at 24hours interval for the next five days (related polygynous male).

Subsequently female mate choice trials were set up as above. 10-day old related monandrous females were given the choice between related polygynous male and unrelated monogynous male. Similar choice was given to related polyandrous females. In corresponding male mate choice trials, both 10-day old related monogynous and polygynous males were given the choice of related monandrous and unrelated polyandrous females. Marking of the adults was done as above. All the trials were conducted in 15 replicates.

### Effect of mating system and relatedness on mating and reproductive parameters

The effect of male and female mating system was observed on following combinations: (1) monandrous female (MA♀)×monogynous male (MG♂), (2) monandrous female (MA♀)×polygynous male (PG♂),(3) monandrous female (MA♀)×related polygynous male (RPG♂), (4) polyandrous female (PA♀)×monogynous male (MG♂), (5) polyandrous female (PA♀)×polygynous male (PG♂), (6) polyandrous female (PA♀)×related polygynous male (RPG♂),(7) monogynous male (MG♂)×polyandrous female (PA♀),(8) monogynous male (MG♂)×related polyandrous female (RPA♀),(8) polygynous male(PG♂)×monandrous female (MA♀),and (9) polygynous male(PG♂)×related monandrous female (RMA♀). Each treatment consisted of 15 replicates. These mating treatments include the pairs obtained from the above mate choice trials. The mating parameters, *i*.*e*. time of commencement of mating (from the introduction in Petri dish to the establishment of genital contact) and copulation duration (from genital contact till the natural disengagement) of the above pairs were recorded. Daily oviposition and percent viability were recorded for the next 7 days in all treatments.

## Data analysis

Data on mate choice was subjected to chi-square (χ^2^) goodness-of-fit analysis. Data on mating parameters and reproductive parameters were first tested for normality (Kolmogorov-Smirnoff) and homogeneous (Bartlett’s) distribution. On being found normally distributed with homogeneous variation, the data were subjected to two-way analysis of variance (ANOVA) with mating system of male and female as individual independent factors. The analysis was followed by the comparison of means using post hoc Tukey’s honest significance test at 5%. All statistical analyses were conducted using R studio Version 1.2.1335 statistical software.

## RESULTS

### Effect of mating system and relatedness on mate choice

Results on mate choice revealed that when monandrous females were given the choice between monogynous and polygynous males then latter were preferred (Fig.1a). Same results were observed when polyandrous females were given the same choice (Fig. 1b). However, when both females were given the choice of related polygynous males and unrelated monogynous males then unrelated monogynous males were preferred as mates (Fig. 1 a and b).

**Figure 1:**
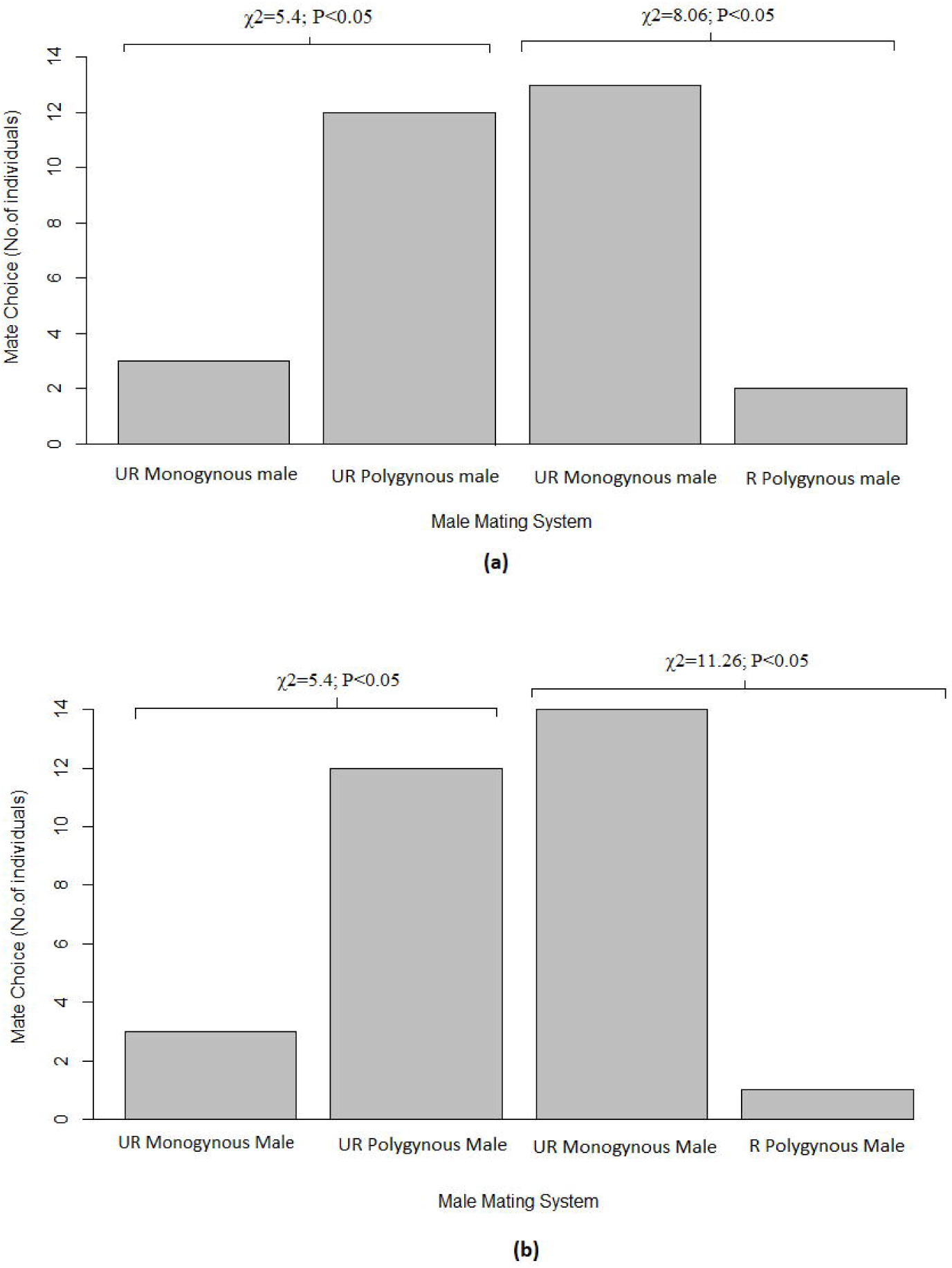
Mate preference of a) monoandrous females, and b) polyandrous females.

In monogynous male mate choice trials, monandrous females were preferred over the polyandrous females (Fig. 2a). Similar choice was made by polygynous males (Fig. 2b). When related monandrous and unrelated polyandrous females were kept in mate choice trials then both males preferred unrelated polyandrous females over the related monandrous females (Fig. 2a and b).

**Figure 2:**
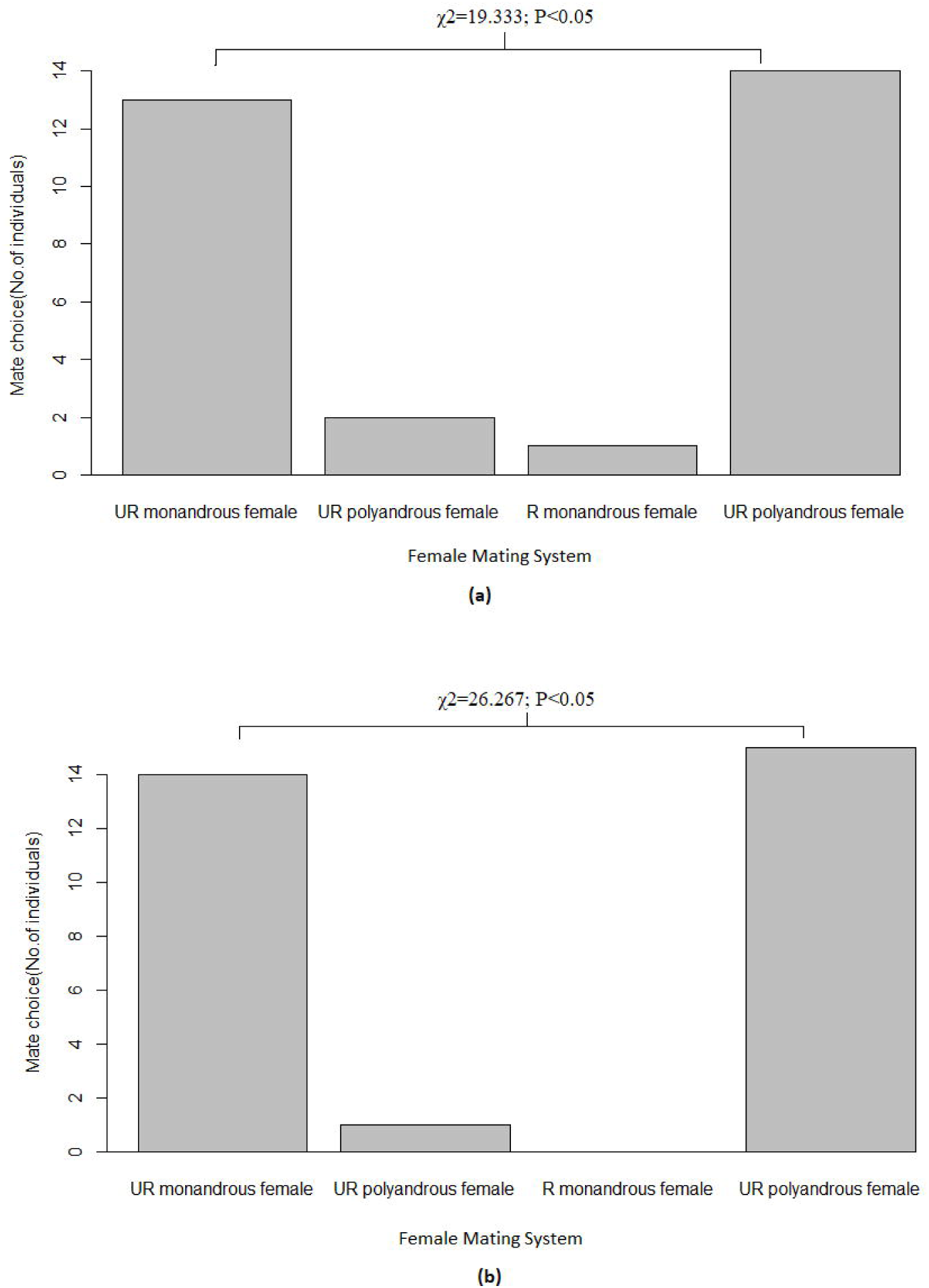
Mate preference of a) monogynous males, and b) polygynous males.

### Effect of mating system and relatedness on mating and reproductive parameters

Time of commencement of mating (TCM) was significantly influenced by the mating system of males (*F*_2, 84_=272.98; *P*=0.000) but not by mating system of females (*F*_2,84_=0.95; *P*=0.332). The interaction between mating system of males and females was also significant (*F*_2, 84_=4.43; *P*=0.014). Time of commencement of mating was longer when monandrous and polyandrous females mated with unrelated monogynous males, followed by related polygynous males, and shortest when mated with unrelated polygynous males (Fig.3a).

When monogynous and polygynous males were mated with females of varying mating system, Time of commencement of mating was not significantly affected by mating system of males (*F*_2,84_=0.02; *P*=0.891) but was significantly influenced by mating system of females (*F*_2,89_=405.72; *P*=0.000) The interaction between mating system of males and females was also significant (*F*_2, 84_=3.93; *P*=0.023). TCM was high when monogynous and polygynous males mated with unrelated polyandrous females and related monandrous females while low when both were mated with unrelated monandrous females (Fig 3b).

**Figure 3:**
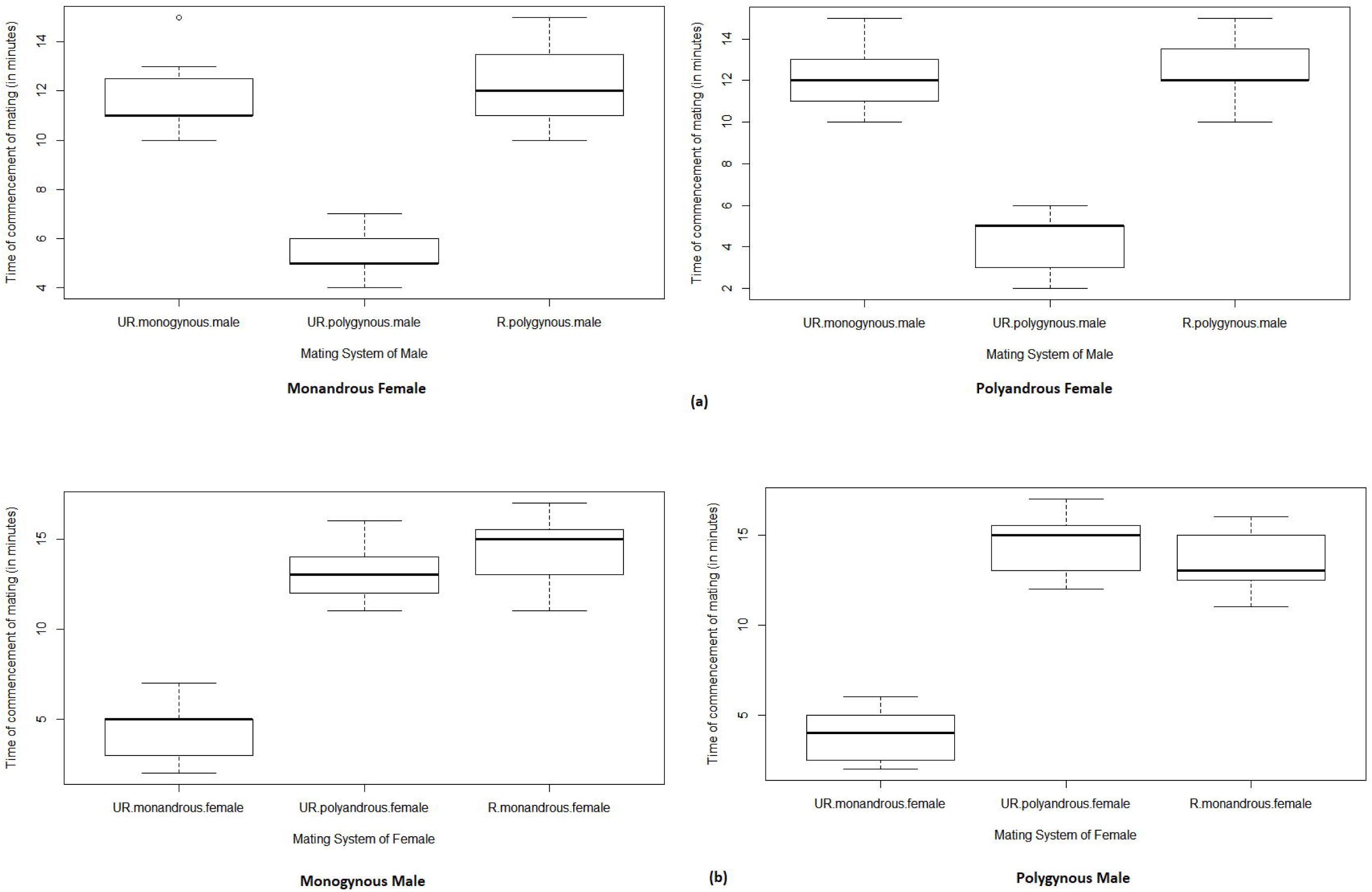
Time of commencement of mating when a) monandrous and polyandrous females, and b) monogynous and polygynous males mated with adults of different mating pattern.

Results of two-way ANOVA revealed the significant effect of male mating system (*F*_2,84_=20.36; *P*=0.000) and insignificant effect of female mating system (*F*_2,84_=2,39; *P*=0.126) on copulation duration when monandrous and polyandrous females mated with males of varying mating system. The interaction between males and females mating system was also insignificant (*F*_2,84_=0.06; *P*=0.937). Copulation duration was highest when monandrous and polyandrous females were mated with unrelated polygynous males and lowest when mated with unrelated monogynous males and related polygynous males (Fig 4a). When males mated with females of different mating systems, copulation duration was significantly influenced by mating system of females (*F*_2,89_=110.15; *P*=0.000)but not by the mating system of males (*F*_1,89_=0.00; *P*=1.000). The interaction between male and female mating systems was also insignificant (*F*_2, 89_=0.43; *P*=0.655). Copulation duration was highest when monogynous and polygynous males mated with unrelated monandrous females (Fig. 4b) and lowest when mated with unrelated polyandrous and related monandrous females (Fig. 4b).

**Figure 4:**
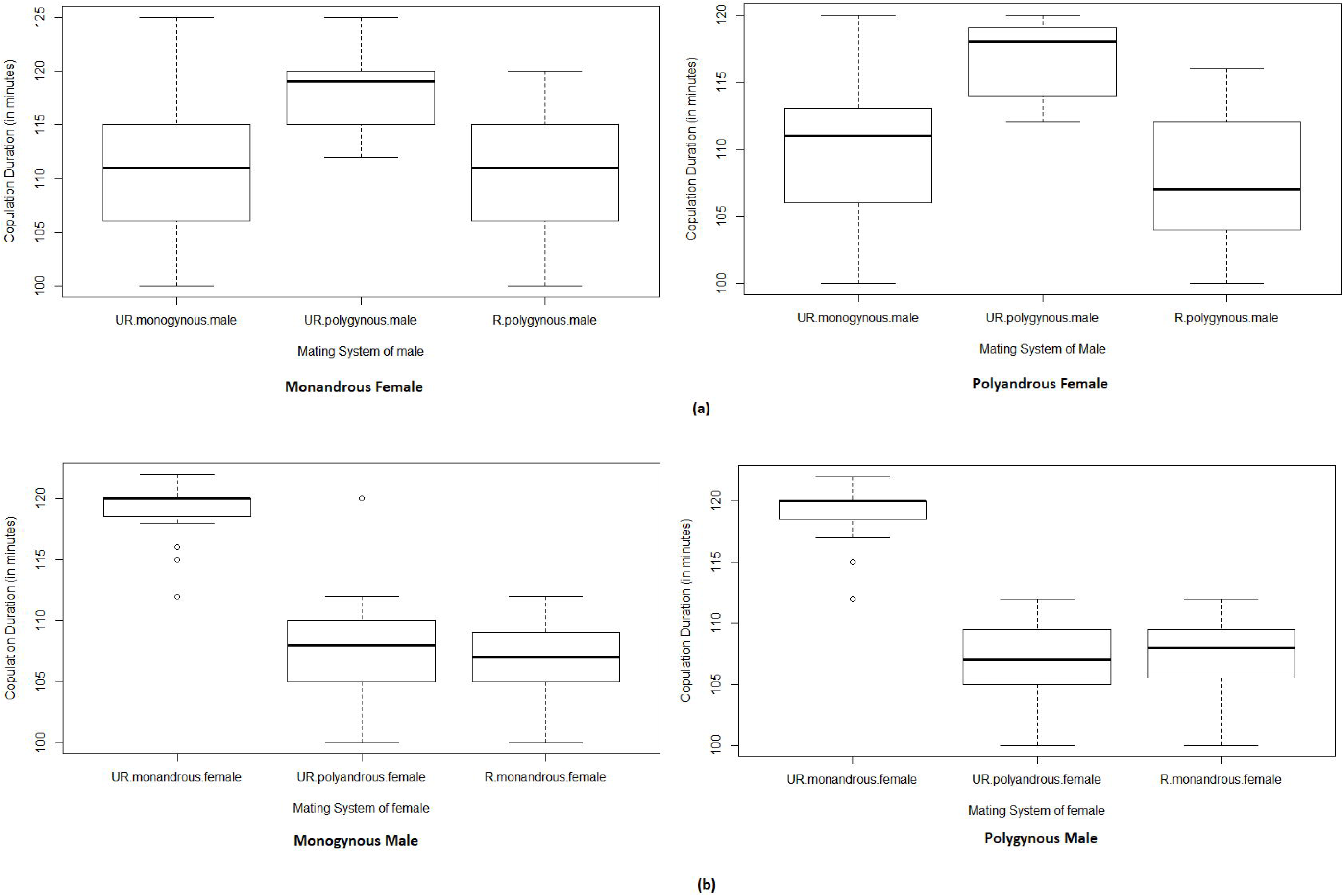
Copulation duration when a)monandrous and polyandrous females, and b) monogynous and polygynous males mated with adults of different mating pattern.

Mating system of males significantly affected fecundity (*F*_2, 89_=2849.94; *P*=0.000) while mating system of females had no significant effect (*F*_1, 89_=0.28; *P*=0.599) when monandrous and polyandrous females were mated with males of varying mating systems. The interaction between mating system of males and females was also insignificant (*F*_2, 89_=0.10; *P*=0.906) Both monandrous and polyandrous females laid more eggs when mated with unrelated polygynous males than when mated with unrelated monogynous males and related polygynous males (Fig. 5a).

**Figure 5:**
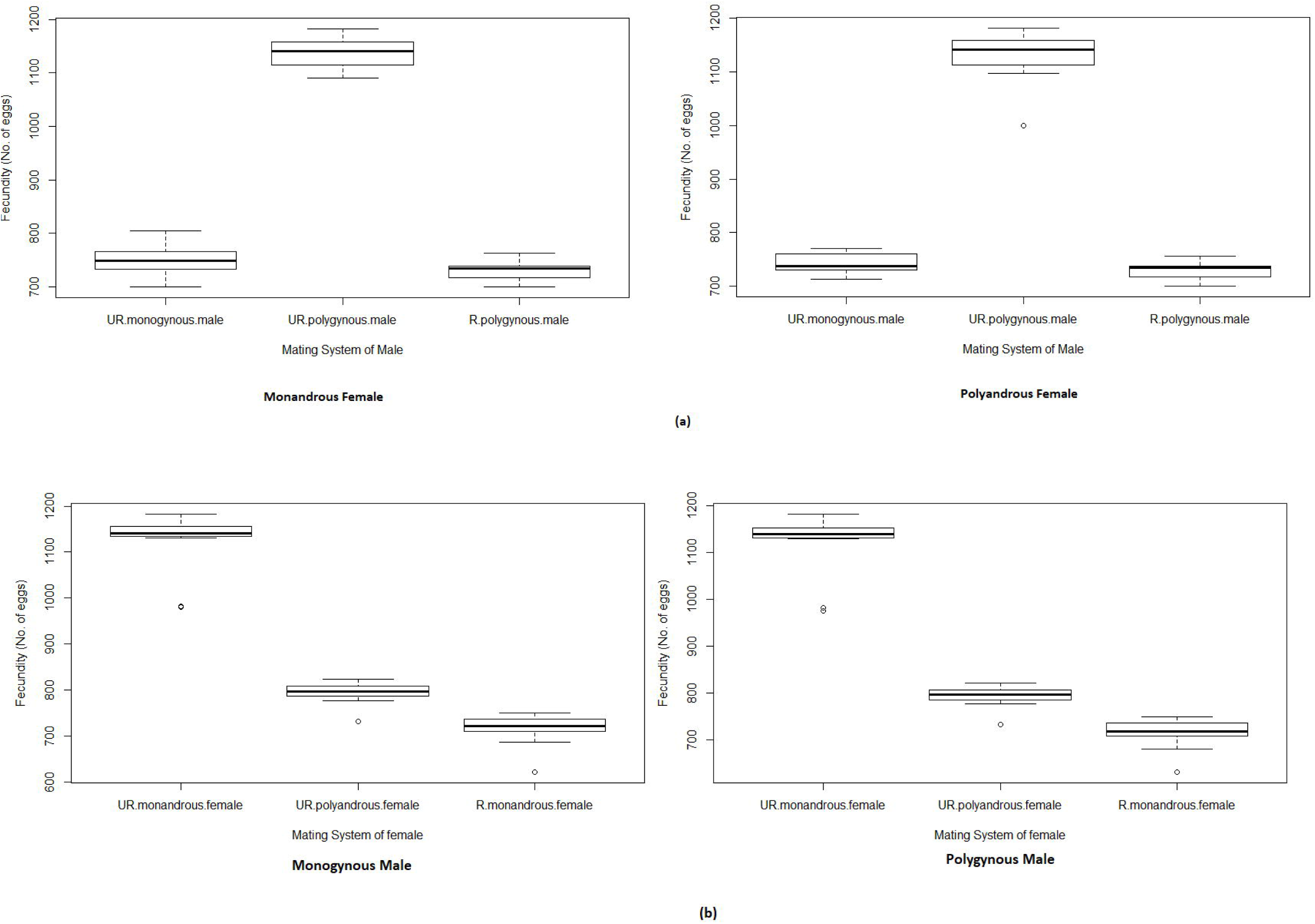
Fecundity when a) monandrous and polyandrous females, and b) monogynous and polygynous males mated with adults of different mating pattern.

When monogynous and polygynous males were mated with females of varying mating system, fecundity was significantly influenced by the mating system of females (*F*_2,89_=3587.51; *P*=0.000) and had no significant effect of the mating system of males (*F*_1,89_=0.00; *P*=0.975) The interaction between males and females mating system was also insignificant (*F*_2,89_=0.07; *P*=0.933). Maximum number of eggs were laid when monogynous and polygynous males mated with unrelated monandrous females (Fig. 5b) and minimum when mated with unrelated polyndrous females and related monandrous females (Fig. 5b).

Percent egg viability was significantly influenced by the mating system of females (*F*_1, 89_=36.31; *P*=0.000) and males (*F*_2,89_=74.91; *P*=0.000) when monandrous and polyandrous females mated with males of varying mating system. The interaction between mating system of males and females was also significant (*F*_2,89_=8.81; *P*=0.000) Percent egg viability was maximum when monandrous and polyandrous females mated with unrelated polygynous males and minimum when mated with related polygynous males (Fig. 6a).

**Figure 6:**
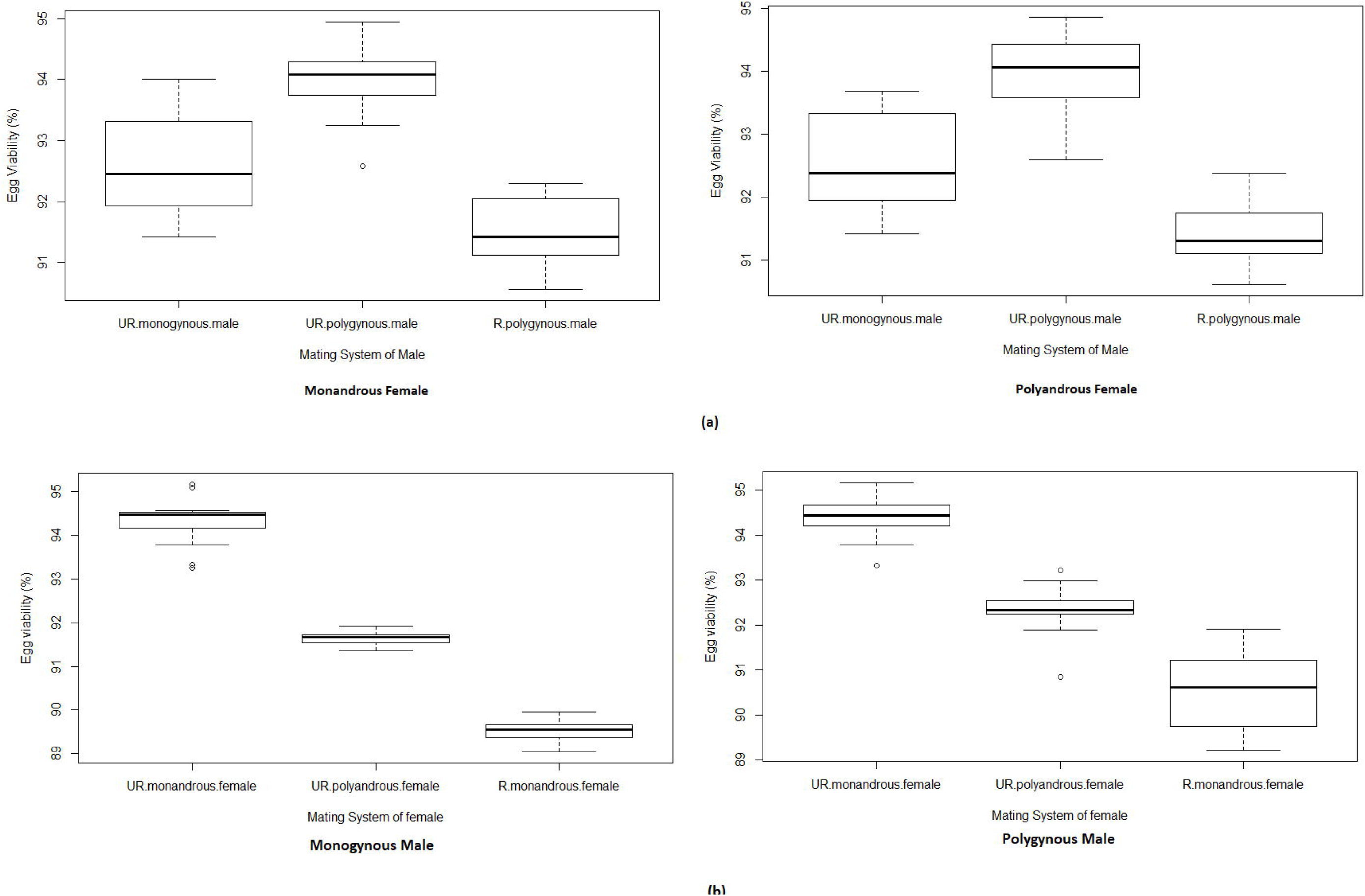
Percent egg viability when a) monandrous and polyandrous females, and b) monogynous and polygynous males mated with adults of different mating pattern.

When monogynous and polygynous males mated with females of varying mating system, both male (*F*_1,89_=6.96; *P*=0.01) and female mating systems (*F*_2, 89_=205.30; *P*=0.000) influenced percent egg viability significantly. The interaction between mating system of males and females was significant (*F*_2,89_=5.79; *P*=0.004) Percent egg viability was recorded more when monogynous and polygynous males mated with unrelated monandrous females and less when both males mated with related monandrous females (Fig. 6b).

## DISCUSSION

The present study reveals that mating system of the mates significantly affects mate choice, mating and reproductive parameters. Monandrous and polyandrous females preferred polygynous males over monogynous one. However, when choice was given between unrelated monogynous and related polygynous males, then unrelated monogynous males were preferred. In male mate choice trials, both monogynous and polygynous males preferred monandrous females over polyandrous females. However, this preference was strongly affected by relatedness, with the choice being reversed if polyandrous females were unrelated and monogamous ones were related.

The preference for polygynous males over monogynous males can be indicative of the fitness of males in terms of more sperm supply and their viability. In *T. casteneum* it was reported that females respond better when they were subjected to more sexually conflicted males (Michalczyk *et al*., 2011). According to the ‘good gene’ hypothesis superior quality males were preferred for direct and indirect benefits (Simmons, 2001). Another reason for preferring the polygynous males is their ability to compete. According to Kvarnemo and Simmons (2013), females prefer to mate with males which are superior competitors and have good genetic quality so that their sons could also have the superior ability to compete.

The biasness for polygynous males was also observed in mating and reproductive parameters. Females mated faster with polygynous males and also mated for the longer duration which can be attributed to their polyandrous nature. Fecundity and percent egg viability was much higher when mated with polygynous males. Studies by Demont et al. (2014) on flour beetle reported that polygynous males are more efficient in terms of providing healthy sperms and get chance to mate more easily. Another reason which can be indicative for the preference of polygynous males is novelty. It has been reported in many studies that new males were preferred over old ones (Madsen et al.,1992; Olsson & Madsen, 1995). Theoretical studies have revealed that mating with multiple mates or new males can benefit their offspring (Yasui, 1998). In bumble bee, *Bombus terrestris* L., it has been reported that mating with different males can reduce the chances of parasitic infection (Baer and Hempel, 1999). Increased number of eggs can be attributed as one of the benefits of polyandry. Arnqvist & Nilsson (2000) reviewed that mating multiple times and with polygynous males have benefits beyond providing sperms, such as more accessory proteins which also enhance the reproductive output of females (Chen, 1984; Gillott, 1988; Eberhard & Cordero, 1995; Eberhard, 1996; Klowden, 1999). Females of *D*.*melanogaster*, due to their polyandrous nature can allocate male sperms and produce more number of eggs (Childress & Hartle, 1972). In ladybirds, reproductive benefits of mating with polygynous males have been reported in *A. bipunctata, Propylea quatuordecimpunctata* (L.), *Harmonia quadripunctata* (Pontoppidan) (Majerus, 1994), *M. sexmaculatus* (F.) (Bind, 2007), *C. septempunctata* (Omkar & Srivastava, 2002), *C. transversalis* (Omkar & James, 2005), *P. dissecta* (Omkar & Mishra, 2005) and *C. saucia* (Omkar et al.,2010).

In male mate choice trials, both monogynous and polygynous males preferred monandrous females. This could possibly be to avoid sperm competition in order to gain more paternity share. Short time to commence mating and longer copulation duration of these pairs can also be attributed to the mate guarding by not allowing their partners to mate with others. More fecundity and percent egg viability could be indicative of increased sperm supply. Males allocate more sperm to virgin females (Engqvist & Reinhold, 2006) or in the presence of competitors (Kelly & Jennions, 2011). Another reason for this preference could be the ‘Coolidge effect’ where males prefer to mate with the novel females (Wilson et al.,1963). In broad-horned beetle, *G. cornutus*, it was reported that males can recognise the cuticular hydrocarbons of the other males and increase their duration of mating as well as their ability to compete with sperms of other males (Lane et al.,2015). Females of many species mating with multiple partners can affect female fecundity or the potential for sperm competition (Carazo et al.,2004). Lane et al. (2015) also reported that males adjust their pre and postcopulatory investments depending on the mating status of the females. Results of various studies also reveals that males use chemical cues to assess the mating status of females (Carazo et al. 2004; Friberg 2006) and risk of sperm competition (Friberg 2006; Carazo et al., 2007; Thomas and Simmons 2009; Garbaczewska et al., 2013). In ladybird *M. sexmaculatus*, also it was reported that males adapted to various strategies such as mate guarding to avoid the risk of sperm competition (Chaudhary et al.,2015) and also preferred unmated females over multiple mated females (Dubey et al., 2018).

In second experiment when relatedness was also included, monandrous and polyandrous females preferred unrelated monogynous males. This could be attributed to inbreeding avoidance. Reduced number of eggs following matings between relatives has also been reported in *D. melanogaster* (Robinson et al., 2012). It has been considered as a postcopulatory strategy for inbreeding avoidance in order to lessen the cost of mating with close relatives (Tregenza & Wedell, 2000). Decreased percent egg viability could be attributed to the biased nature of females for the sperms of related males (Charlesworth & Charlesworth, 1987; Zeh & Zeh, 1996, 1997; Tregenza & Wedell, 2000) and also the poor offspring quality (Keller &Waller, 2002). In *M. sexmaculatus* it was established that related individuals are less preferred over unrelated individuals (Saxena et al.,2016).

Thus, it can be concluded from the above study that (i) both monandrous and polyandrous females preferred polygynous males over monogynous in order to choose a broader gene pool for their offspring, (ii) between related polygynous and unrelated monogynous males, unrelated males were preferred to avoid inbreeding (iii) when monogynous and polygynous males were subjected to mate choice trials both prefer monandrous females in order to reduce the risk of sperm competition.

## Acknowledgements

SS also acknowledges BSR Fellowship by University Grant Commission, New Delhi, India (F.No.25-1/2014-15 (BSR)/7-109/2007/BSR) dated August 25, 2015.

## Figure Legends

Figure 1: Mate preference of a) monoandrous females, and b) polyandrous females.

Figure 2: Mate preference of a) monogynous males, and b) polygynous males.

Figure 3: Time of commencement of mating when a) monandrous and polyandrous females, and b) monogynous and polygynous males mated with adults of different mating pattern.

Figure 4: Copulation duration when a)monandrous and polyandrous females, and b) monogynous and polygynous males mated with adults of different mating pattern.

Figure 5: Fecundity when a) monandrous and polyandrous females, and b) monogynous and polygynous males mated with adults of different mating pattern.

Figure 6: Percent egg viability when a) monandrous and polyandrous females, and b) monogynous and polygynous males mated with adults of different mating pattern.

